# Unlabeled salivary gland organoids have distinct Raman signatures following FGF2-induced proacinar cell differentiation

**DOI:** 10.1101/2021.09.16.460651

**Authors:** Kate Tubbesing, Nicholas Moskwa, Ting Chean Khoo, Deirdre A. Nelson, Anna Sharikova, Melinda Larsen, Alexander Khmaladze

## Abstract

Organoids are self-organized three-dimensional (3D) tissue cultures that model the structure and function of organs to provide insights into signaling during organ formation and have translational applications in disease modeling and assessing drug responses. Due to their heterogeneity, there is a need for non-destructive methods to identify the differentiation state, or the phenotype, of organoids. As organoids often contain complex mixtures of basement membrane and/or extracellular matrix proteins, which are often highly auto-fluorescent, it typically makes low-resolution Raman measurements a challenge. We developed Raman confocal micro-spectroscopy methods to avoid and minimize the matrix signal and define specific Raman signatures for growth factor-differentiated and non-differentiated organoids. In complex, branched salivary gland organoids derived from mouse embryonic epithelial and stromal cells embedded within the laminin-rich basement membrane matrix, Matrigel, we identified specific Raman spectral signatures for organoids in different differentiation states. We report that either comparison of spectral signatures or multivariate SVD analysis can be used to distinguish between organoids treated with FGF2, organoids treated with EGF, and non-treated controls. Raman spectral signatures can be used to non-invasively distinguish between different phenotypes in the 3D context of unlabeled organoids.

**Highlights:**

- FGF2-dependent proacinar cell differentiation in salivary organoids have unique Raman signatures detected with a novel confocal-based Raman imaging approach.
- These signatures can be used in unlabeled salivary organoids to monitor proacinar cell differentiation.
- Confocal-based Raman imaging may be applicable to monitoring differentiation state of other types of organoids.

## Introduction

Organoids are essential biological tools, utilized across multiple fields, as they better represent the 3D organization and function of multiple cell types and often provide translationally valuable insight into drug response, disease pathology, and developmental biology [1]. In general, organoids are created through the co-culture of multiple types of either embryonic or stem cells, within an extracellular 3D matrix environment [1,2]. The correct combination of cell types and media, which is often modified to promote developmental signaling, will result in the 3D organization of cells with distinct organ-like structures. There are various protocols to produce organoids; all of them are labor, material, and time intensive. Organoids that express fluorescent or luciferase proteins have numerous screening methods [3–5], however the use of protein expression systems is not well suited for all organoid models and pre-clinical applications. Therefore, there is a need for non-destructive methods to confirm developmental or phenotypic stages of organoids without fluorescent labeling.

Raman imaging methods vary in resolution, but in general offer a non-destructive, label-free approach to defining biological samples [6]. The Raman effect is a natural phenomenon of inelastic light scattering, which is determined by the vibrational energy levels of specific molecular structures [7]. The interpretation of Raman spectra of biological samples is often dependent on the spatial resolution of the method. In clinical settings, low-resolution fiber-based Raman measurements allow the discrimination of healthy tissue regions from disease- or tumor-burdened regions [8,9]. The higher resolution approaches implemented with Raman micro-spectroscopy can be used to gain information on sub-cellular or cellular populations, with the scale determined by the numerical aperture of the microscope objective and the confocal settings of the instrument [10–13]. Importantly, while Raman spectroscopy is often compared to Fourier transform infrared (FTIR) spectroscopy, the hydrated biological samples are better suited for Raman measurements that do not have the interference from water [14].

As organoids contain complex mixtures of basement membrane or extracellular matrix (ECM) proteins, which are often highly auto-fluorescent, it typically makes low-resolution Raman measurements a challenge. To minimize this issue, we developed methods to minimize the ECM signal and use Raman confocal micro-spectroscopy to define specific Raman signatures of growth factor-differentiated and non-differentiated organoids. The salivary gland organoid model utilized to validate the Raman approach is well characterized and consists of epithelial and stromal cells derived from mouse embryonic salivary glands embedded within the laminin-rich basement membrane matrix Matrigel [2,15]. Salivary glands consist of branching ductal systems with terminal buds of saliva producing acinar epithelial cells. The salivary gland organoid model utilized throughout produces the desired proacinar and ductal regions with FGF2 treatment, while untreated or EGF-treated organoids lack the differentiated proacinar cells [2,15]. Development of this Raman-based organoid imaging method has broad application to numerous organoid and 3D culture models.

## Results

### FGF2 promotes proacinar and ductal phenotype differentiation in salivary gland organoids

Using our established protocol, the salivary gland proacinar organoids were formed from embryonic epithelial and stromal cells and were treated with FGF2, EGF, or control media lacking growth factor (**Figure 1A**). Organoids treated with FGF2, but not with EGF, or no additional growth factor supplementation, exhibited robust proacinar cell differentiation with buds expressing membrane-localized aquaporin-5 (AQP5) (**Figure 1B**). Immunofluorescent antibody staining was performed to recognize epithelial cells expressing aquaporin-5 to identify proacinar/acinar cells, and keratin-7 (K7) to identify ductal cells. As expected, the AQP5/DAPI ratio was significantly increased with FGF2 treatment (**Figure 1C)**. To determine whether FGF2 induced a change in phenotype of the epithelial cells, immunostaining was performed to detect the ductal protein, cytokeratin 7 (K7). The enrichment of ductal cells (K7/DAPI ratio) was observed in the control, but not in the EGF- or FGF2-treated cells (**Figure 1D**). The ratio of the image area of proacinar cells (AQP5) to ductal cells (K7) was significantly increased in FGF2-treated organoids (**Figure 1E**). These data show that the FGF2-, EGF-treated, and control organoids represent three different epithelial differentiation states.

**Figure 1.**
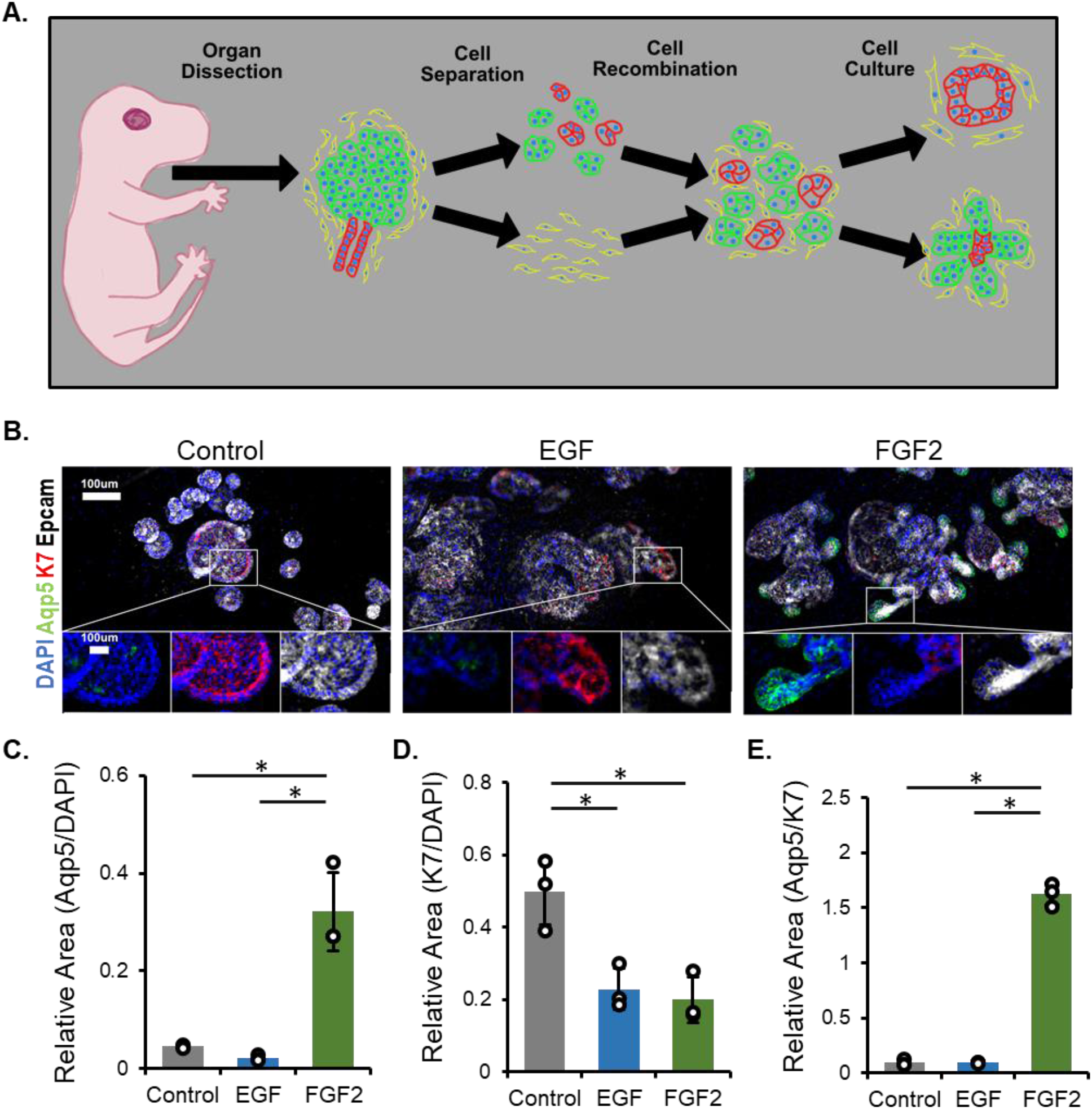
FGF2 promotes pro-acinar and inhibits ductal differentiation in embryonic salivary gland organoids. (**A**) Illustration of salivary gland organoid creation from murine embryonic salivary gland cells, which are separated for regulated recombination in Matrigel. Keratin-7 (K7) positive ductal cells are shown in red, aquaporin-5 (AQP5) positive pro-acinar cells are shown in green, and stromal mesenchymal cells are shown in yellow. (**B**) Representative images show that while all organoids express the pan-epithelial marker EpCAM (white) and some level of the ductal marker K7 (red), the elaboration of AQP5+ cells (green) is restricted to branched organoids treated with FGF2. Nuclei are stained with DAPI. (**C-E**) Quantification of microscopy images with relative area ratios for AQP5 to DAPI, K7 to DAPI and AQP5 to K7. Error bar represents 95% confidence interval, n=3 technical replicates, asterisk indicates significance with p>0.05 (Anova-Tukey) (statistical summary in **Supplementary Table 1-3**).

### Development of Raman confocal method to evaluate salivary gland organoids

A significant complication in Raman spectroscopy of organoids involves mitigation of autofluorescence from Matrigel and similar 3D matrix or scaffolding materials. Here, a method employing confocal Raman micro-spectroscopy was developed to identify the cell dense regions of the organoids. To avoid fluorescence from plastic dishes, the unlabeled organoids were transferred to a quartz-based chamber, and the organoids remained submerged in PBS for imaging. The average size of the organoid was determined while in the culturing dish, and it was shown to be larger for the growth factor treated organoids, compared to untreated organoids (**Figure 2A**). Upon transfer to the quartz-based chamber, the cell-dense regions, representing regions composed primarily of epithelial cells, were identifiable and distinct from the Matrigel-dense regions, representing stromal cell-rich regions (**Figure 2B**). After the X-Y coordinates of the cell-dense regions were identified using the brightfield imaging, a series of Raman confocal measurements along the Z direction was performed to focus on the cell dense volume and exclude the spectra from the highly-fluorescent Matrigel. The laser and acquisition settings were selected to facilitate the easy separation of Matrigel-rich regions from cell dense regions (**Figure 2C**). Further, to reduce human error and facilitate high-throughput processing, we confirmed that multivariate SVD analysis can be used to distinguish the Matrigel spectra from the epithelial cell-derived spectra (**Figure 2C**).

**Figure 2.**
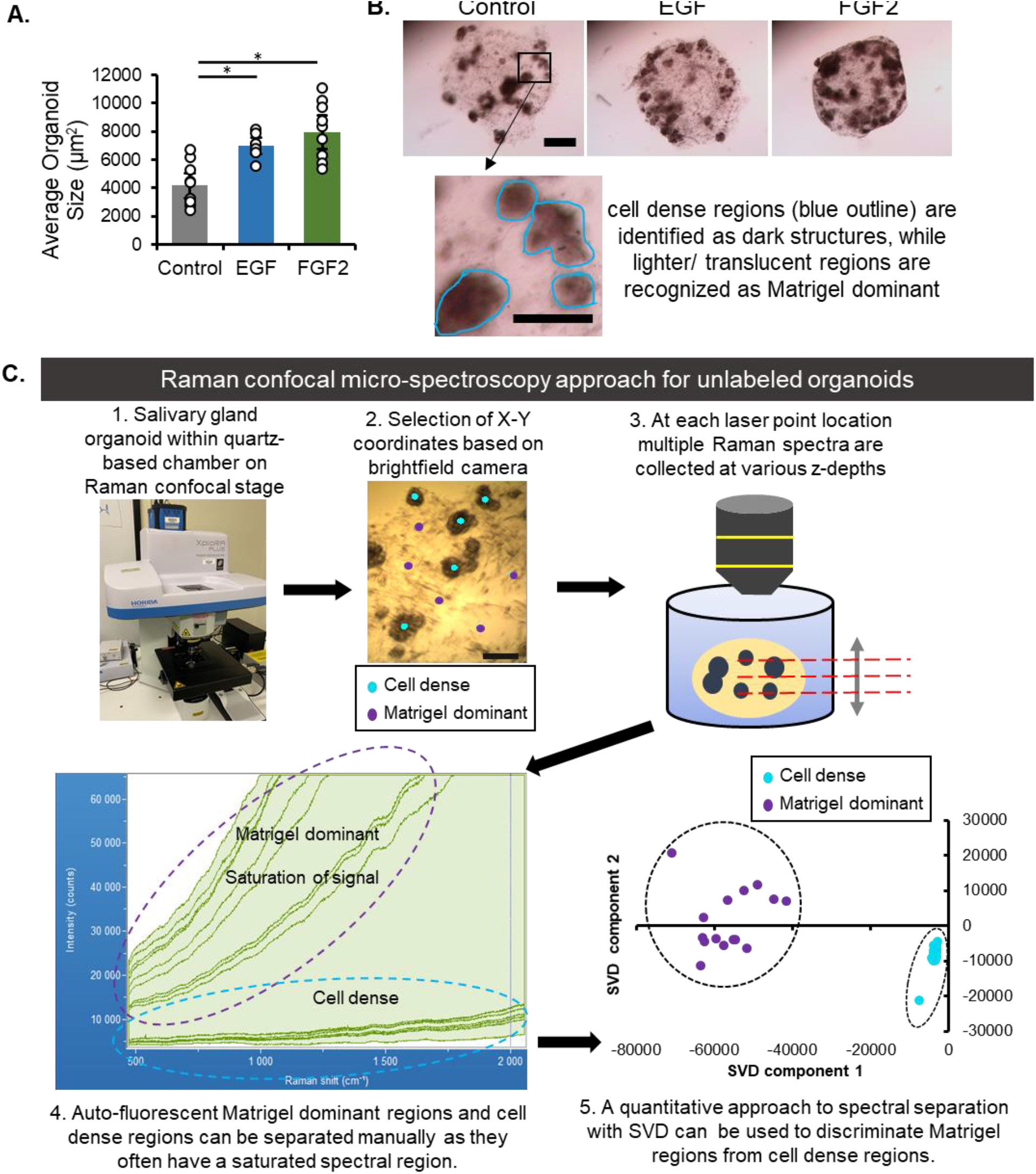
Development of Raman confocal micro-spectroscopy method to evaluate salivary gland organoids. (**A**) Quantification of organoid size based on area. n=7 technical replicates across 3 experiments (statistical summary in **Supplementary Table 4**). (**B**) Representative images of control and growth factor-treated organoids prior to Raman imaging, with the cell-dense, largely epithelial regions highlighted in zoomed panel. Scale bar 500 µm (panel), 250 µm (zoomed region). (**C**) Visualization of the Raman confocal micro-spectroscopy method for identification of cell-dense regions. This method allows the separation of cell dense-organoid regions from Matrigel-dominant, or stromal, regions, which can be done either manually by peak comparison or with a multivariate SVD approach.

### FGF2 treated organoids have a Raman signature which is distinct from untreated or EGF treated organoids

Differentiation state-specific spectra were determined following the removal of the Matrigel spectra. Raman measurements were collected from the cell-dense, epithelial enriched regions of the organoids, which appear as dark brown spheres in brightfield images, **Figure 3A**. The average Raman spectra for control (untreated), EGF-treated and FGF2-treated organoids were distinct from each other, with several peaks having significant differences in intensities (**Figure 3B**). Select peak ratios could be used to differentiate at least one organoid treatment from the other two (highlighted in **Figure 3B**, analysis in **Figures 3C-K**). Some peak ratios showed significant differences between all groups (**Figures 3C, 3E, 3F**), while others could only separate out one treatment from the other two (**Figures 3D, 3G-K**). Many of these peaks have previously been assigned to specific compounds and cellular components. Known specific peak assignments are indicated in **Table 1**, and broad peaks in **Supplementary Figure 1**, but it is not the focus of this study. These data demonstrate that there are specific ratios that differ between organoids in different differentiation states.

**Figure 3.**
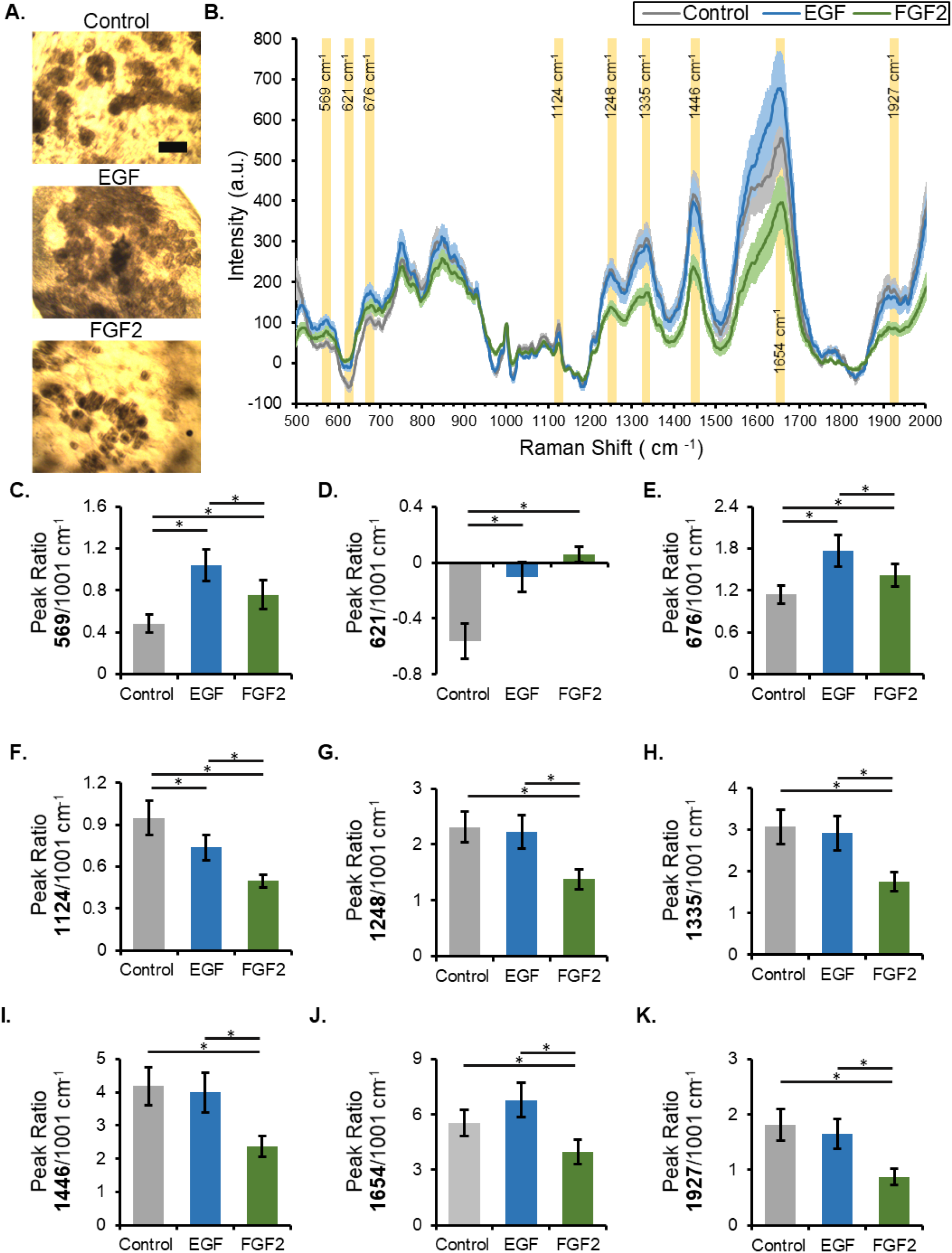
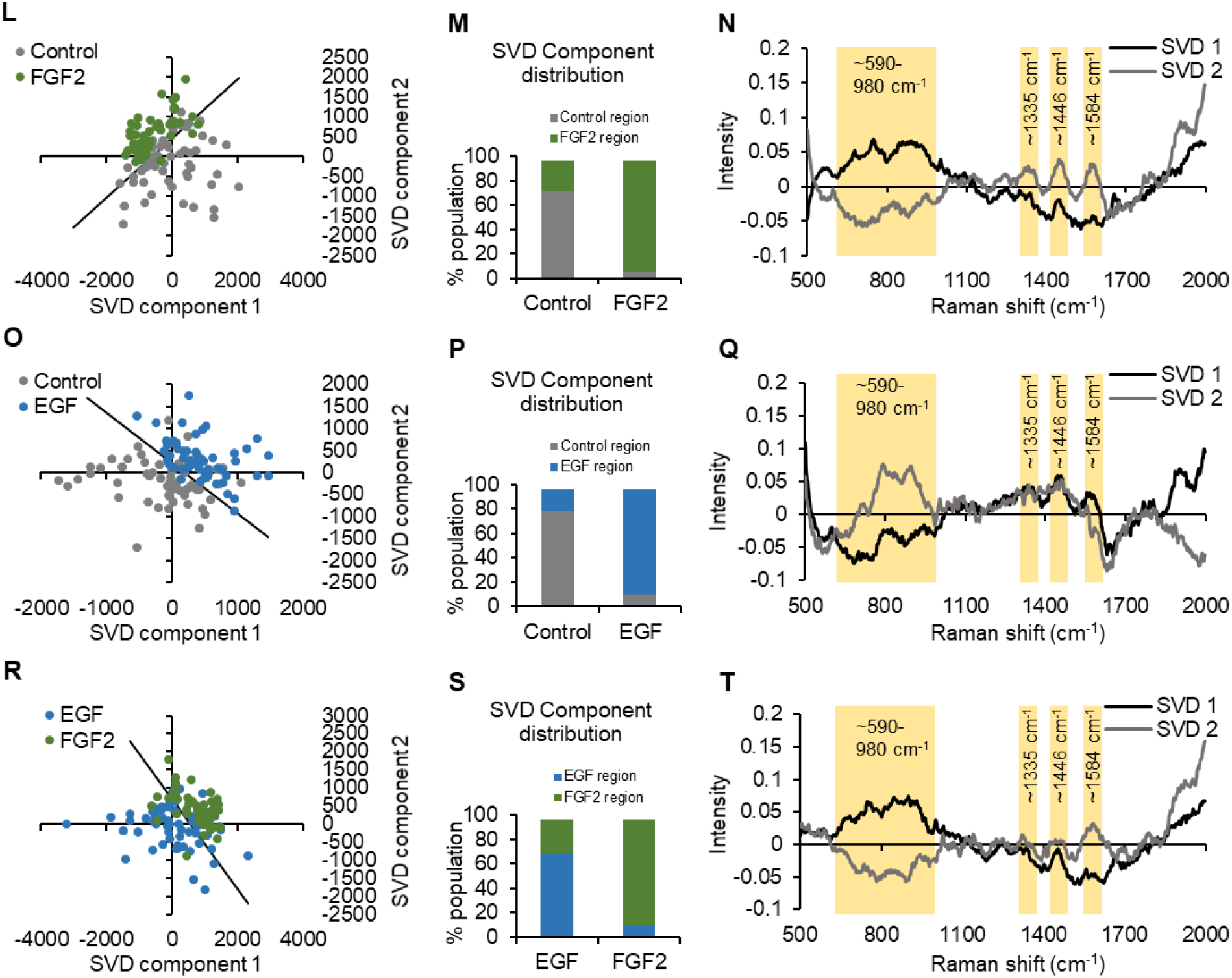
FGF2-treated organoids have a Raman signature which is distinct from EGF-treated or untreated organoids. (**A**) Representative brightfield images of organoids on the Raman confocal stage; the cell-dense regions (dark brown) are easily distinguished from the Matrigel-dominant area (scale bar is 200 µm). (**B**) Average Raman spectra for cell dense regions of control, EGF-, and FGF2-treated organoids, with error bars representing the 95% confidence interval. (**C-K**) Quantification of select peak intensities in the spectral dataset based on the ability to differentiate one or more treatments (n=50 separate spectra collected from a minimum of 3 independent organoids, ^*^ asterisk indicates significance with p<0.05, statistical summary in **Supplementary Table 5**). (**L-T**) Multivariate SVD analysis with control vs. FGF2 (**L-N**), control vs. EGF (**O-P**), and EGF vs. FGF2 (**R-T**). For each comparison there is an SVD plot where each dot represents a single spectrum, with clustering of similar spectra (**L, O, R**). A line in each SVD plot is placed perpendicular and equidistant to the mean of each treatment to separate the treatments in an unbiased manner. The distribution of the spectra in each region is plotted in **M, P, and S**, while the SVD components producing the distribution are plotted in **N, Q, and T**. Select regions of interest within the component graphs are highlighted in yellow (n=50 separate spectra collected from a minimum of 3 independent organoids).

**Table 1.**
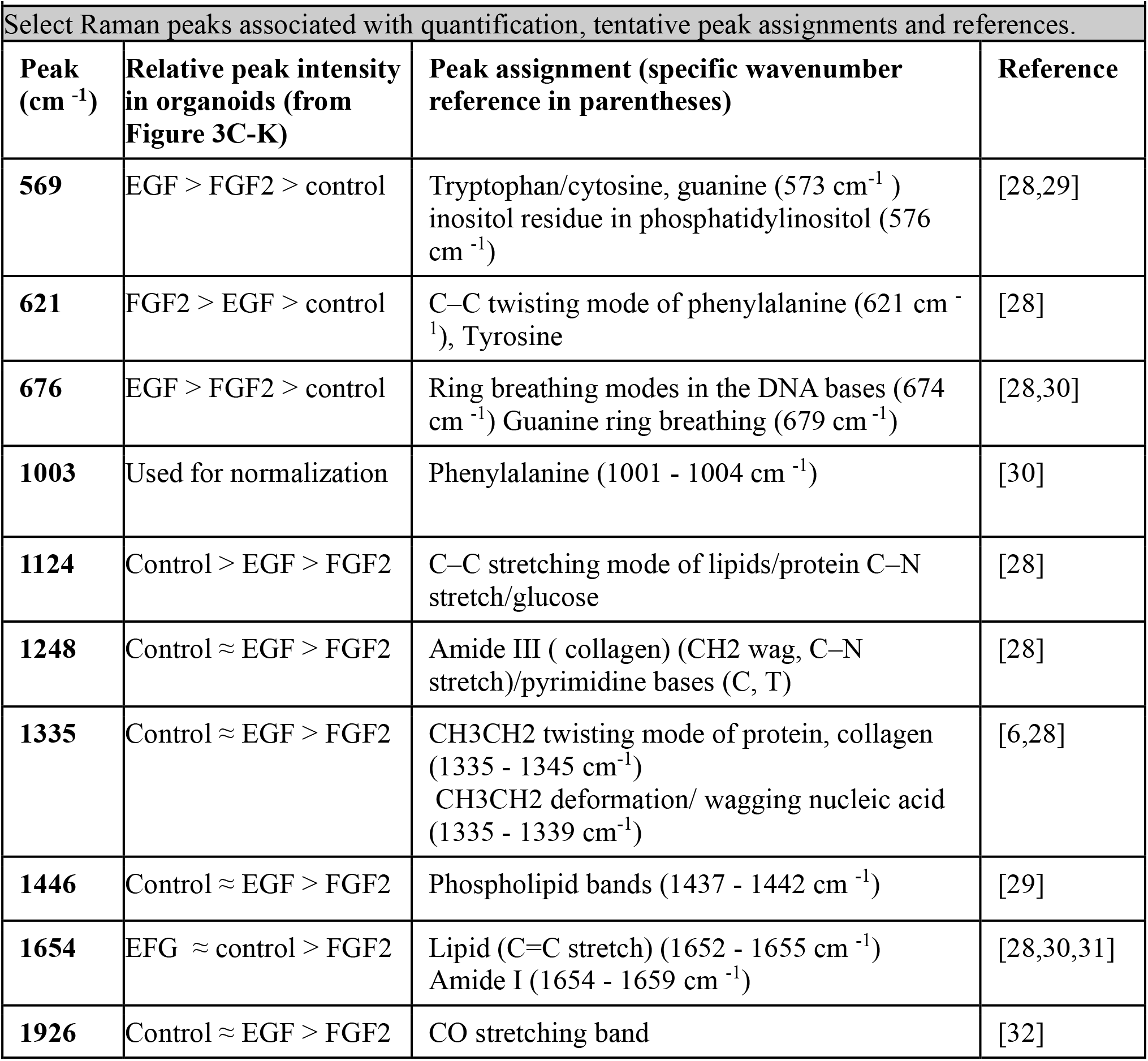
Raman peak assignments reference table.

**Select Raman peaks associated with quantification, tentative peak assignments and references.**
| **Peak (cm<sup>-1</sup>)** | **Relative peak intensity in organoids (from Figure 3C-K)** | **Peak assignment (specific wavenumber reference in parentheses)** | **Reference** |
| --- | --- | --- | --- |
| **569** | EGF > FGF2 > control | Tryptophan/cytosine, guanine (573 cm<sup>-1</sup>)<br>inositol residue in phosphatidylinositol (576 cm<sup>-1</sup>) | [28,29] |
| **621** | FGF2 > EGF > control | C–C twisting mode of phenylalanine (621 cm<sup>-1</sup>), Tyrosine | [28] |
| **676** | EGF > FGF2 > control | Ring breathing modes in the DNA bases (674 cm<sup>-1</sup>) Guanine ring breathing (679 cm<sup>-1</sup>) | [28,30] |
| **1003** | Used for normalization | Phenylalanine (1001 - 1004 cm<sup>-1</sup>) | [30] |
| **1124** | Control > EGF > FGF2 | C–C stretching mode of lipids/protein C–N stretch/glucose | [28] |
| **1248** | Control ≈ EGF > FGF2 | Amide III ( collagen) (CH<sub>2</sub> wag, C–N stretch)/pyrimidine bases (C, T) | [28] |
| **1335** | Control ≈ EGF > FGF2 | CH<sub>3</sub>CH<sub>2</sub> twisting mode of protein, collagen (1335 - 1345 cm<sup>-1</sup>)<br>CH<sub>3</sub>CH<sub>2</sub> deformation/ wagging nucleic acid (1335 - 1339 cm<sup>-1</sup>) | [6,28] |
| **1446** | Control ≈ EGF > FGF2 | Phospholipid bands (1437 - 1442 cm<sup>-1</sup>) | [29] |
| **1654** | EGF ≈ control > FGF2 | Lipid (C=C stretch) (1652 - 1655 cm<sup>-1</sup>)<br>Amide I (1654 - 1659 cm<sup>-1</sup>) | [28,30,31] |
| **1926** | Control ≈ EGF > FGF2 | CO stretching band | [32] |

Multivariate analysis of Raman spectral data was performed in addition to the analysis of individual peak ratios. The entire spectral dataset was subjected to the singular value decomposition (SVD) multivariate analysis (**Figures 3L-T**). The SVD analysis facilitates the unbiased processing of large spectral datasets. The spectra with similar features will cluster together on the SVD scatter plot, while the SVD components provide insight into the specific spectral regions that differentiate the samples [10]. A line systematically placed in each SVD scatter plot (**Figures 3L, 3O & 3R**) is perpendicular and equidistant to the individual mean of each cluster; it is used to quantify the separation of treatments (**Figures 3M, 3P & 3S**). The qualitative analysis shows that, within each comparison made, there is clustering together of the spectra from the same treatment group (**Figure 3L, 3O, 3R)**. Quantification of the data shows that FGF2 and EGF treatments have 10-20% overlap when compared to control **(Figures 3M, 3P)**. The comparison of the two growth factor treatments results in 10-30% of spectra being over the line separating the groups **(Figure 3S**). This is not unexpected, since all samples started with the same cellular material and some ductal cells are present in all treatments, while only proacinar differentiation is observed in the FGF2 treated samples. The leading SVD components share similar trends, as expected, because the broad peaks represent structures which are highly abundant in biological samples (**Figures 3N, 3Q & 3T**); the highlighted regions have notable variations between the treatment groups. Not all peaks that are prominent in the SVD components correspond to the peaks selected for analysis of peak ratios (**Figures 3G-K**). This shows that the multivariate analysis contains additional information that is not limited to several well-defined peaks; for example, the very broad peaks spanning 100 wavenumbers have significant variations between treatments.

### Blinded organoid samples successfully identified with two distinct approaches

To determine whether organoid differentiation state can be predicted on the basis of a Raman signature alone, a blinded study was performed, in which 10 distinct spectra obtained from 3 unknown groups were assigned a control, EGF-treated, or FGF2-treated phenotype (**Figure 4A)**. Two separate approaches were utilized, one based on the multiple-peak comparisons and the other based on the SVD analysis. Multiple-peak comparisons used the mean ratios of Raman peaks previously identified as showing significant differences between organoid phenotypes (**Figures 3B-K**). The unknown sample spectra were averaged and normalized, and their mean ratios were compared to the known Raman peak ratios (**Figures 4B-J**). Following the assignment of individual peaks based on the mean peak ratios, a simple majority was used to classify the unknown samples (**Table 2**). All unknown organoid samples were correctly identified as control, EGF-, or FGF2-treated using multiple Raman peak comparisons.

**Table 2.**
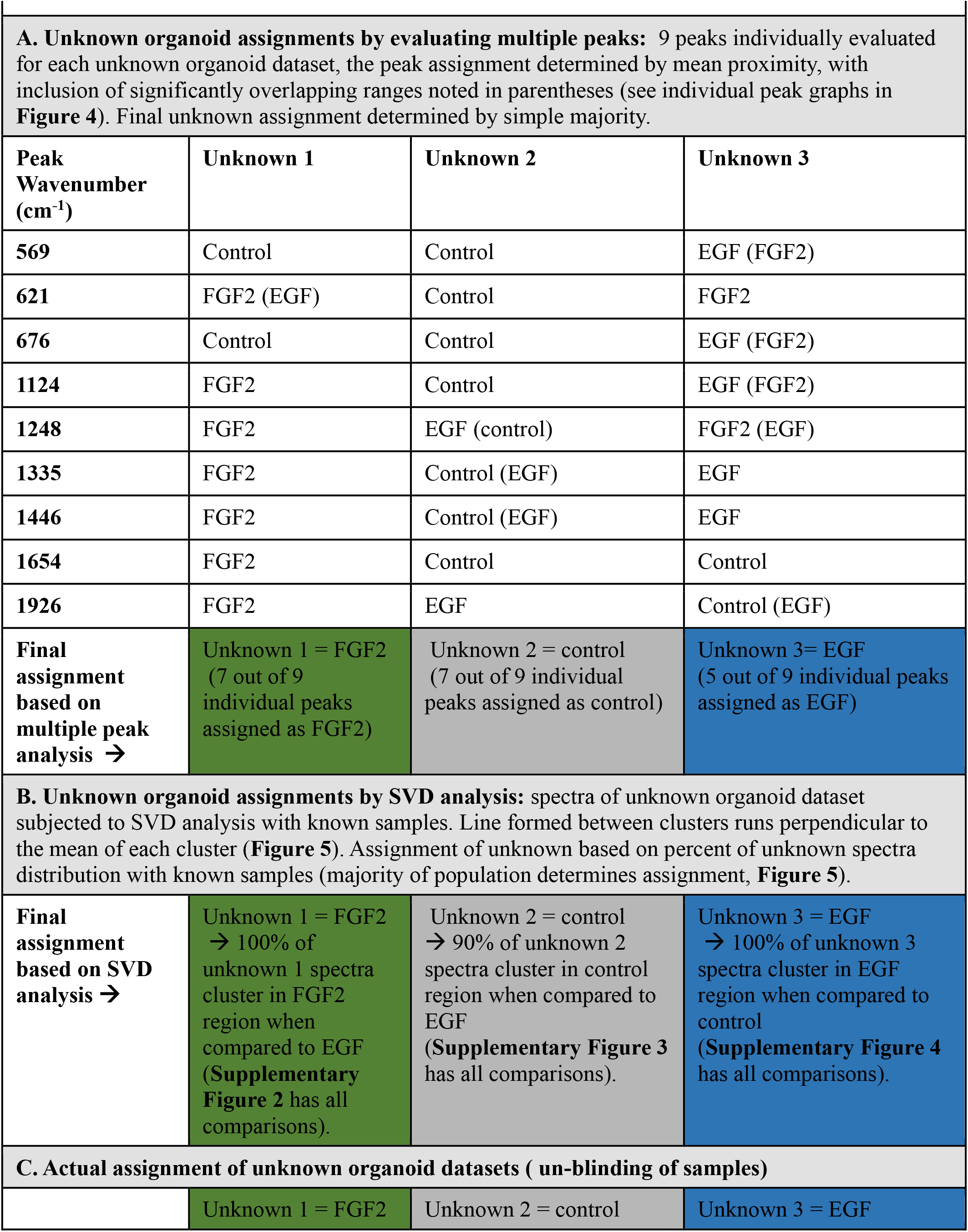
Determination of unknown samples using two distinct approaches.

| Peak Wavenumber (cm<sup>-1</sup>) | Unknown 1 | Unknown 2 | Unknown 3 |
| --- | --- | --- | --- |
| 569 | Control | Control | EGF (FGF2) |
| 621 | FGF2 (EGF) | Control | FGF2 |
| 676 | Control | Control | EGF (FGF2) |
| 1124 | FGF2 | Control | EGF (FGF2) |
| 1248 | FGF2 | EGF (control) | FGF2 (EGF) |
| 1335 | FGF2 | Control (EGF) | EGF |
| 1446 | FGF2 | Control (EGF) | EGF |
| 1654 | FGF2 | Control | Control |
| 1926 | FGF2 | EGF | Control (EGF) |
| **Final assignment based on multiple peak analysis →** | Unknown 1 = FGF2 (7 out of 9 individual peaks assigned as FGF2) | Unknown 2 = control (7 out of 9 individual peaks assigned as control) | Unknown 3 = EGF (5 out of 9 individual peaks assigned as EGF) |
| **Final assignment based on SVD analysis →** | Unknown 1 = FGF2 → 100% of unknown 1 spectra cluster in FGF2 region when compared to EGF (**Supplementary Figure 2** has all comparisons). | Unknown 2 = control → 90% of unknown 2 spectra cluster in control region when compared to EGF (**Supplementary Figure 3** has all comparisons). | Unknown 3 = EGF → 100% of unknown 3 spectra cluster in EGF region when compared to control (**Supplementary Figure 4** has all comparisons). |
| **C. Actual assignment of unknown organoid datasets ( un-blinding of samples)** |  |  |  |
|  | Unknown 1 = FGF2 | Unknown 2 = control | Unknown 3 = EGF |

**Figure 4.**
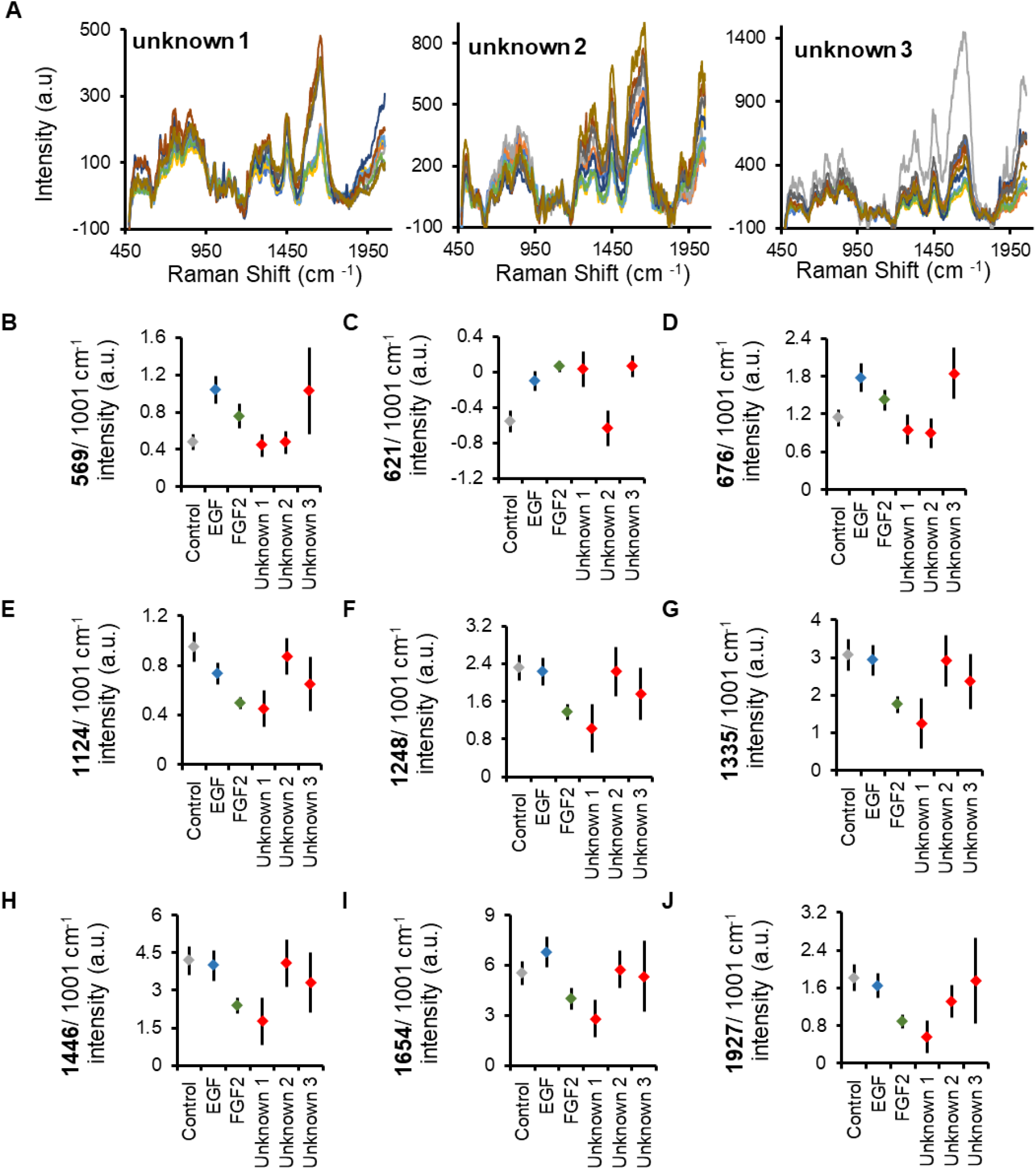
Unknown organoids identified using analysis of select Raman spectral peaks. (**A**) Raman spectra of cell-dense organoid region for each unknown (blinded) organoid dataset (10 processed, normalized spectra per treatment). (**B-J**) Comparison of the mean Raman peaks between the known samples (full spectra in Figure 3) and the unknown data sets. Error bars equal 95% confidence interval. **Table 2** details the assignment of the unknown peaks to specific treatments.

An alternative approach for identifying unknown organoid samples utilized the multivariate SVD analysis that was also used in the evaluation of Raman signatures of organoid phenotypes (**Figures 3L-T**). Each unknown sample, consisting of 10 spectra, was compared individually to the known datasets (two known datasets at a time) until a significant overlap was found (**Figures 5A, 5C, 5E and Supplementary Figures 2-4**). A simple quantification of the unknown sample distribution is easily computed (**Figures 5B, 5D, 5F**). Here it was important to evaluate all comparisons and recognize that the unknown being compared to the wrong datasets could result in 50% overlap (**Supplementary Figures 2-4**). The final assignment of each unknown organoid is included in **Table 2**, and the correct assignment was made with both simple majority and SVD approach to the blinded organoid analysis. All unknown organoid samples were correctly identified as control, EGF-, or FGF2-treated using multivariate SVD analysis of their Raman signals.

**Figure 5.**
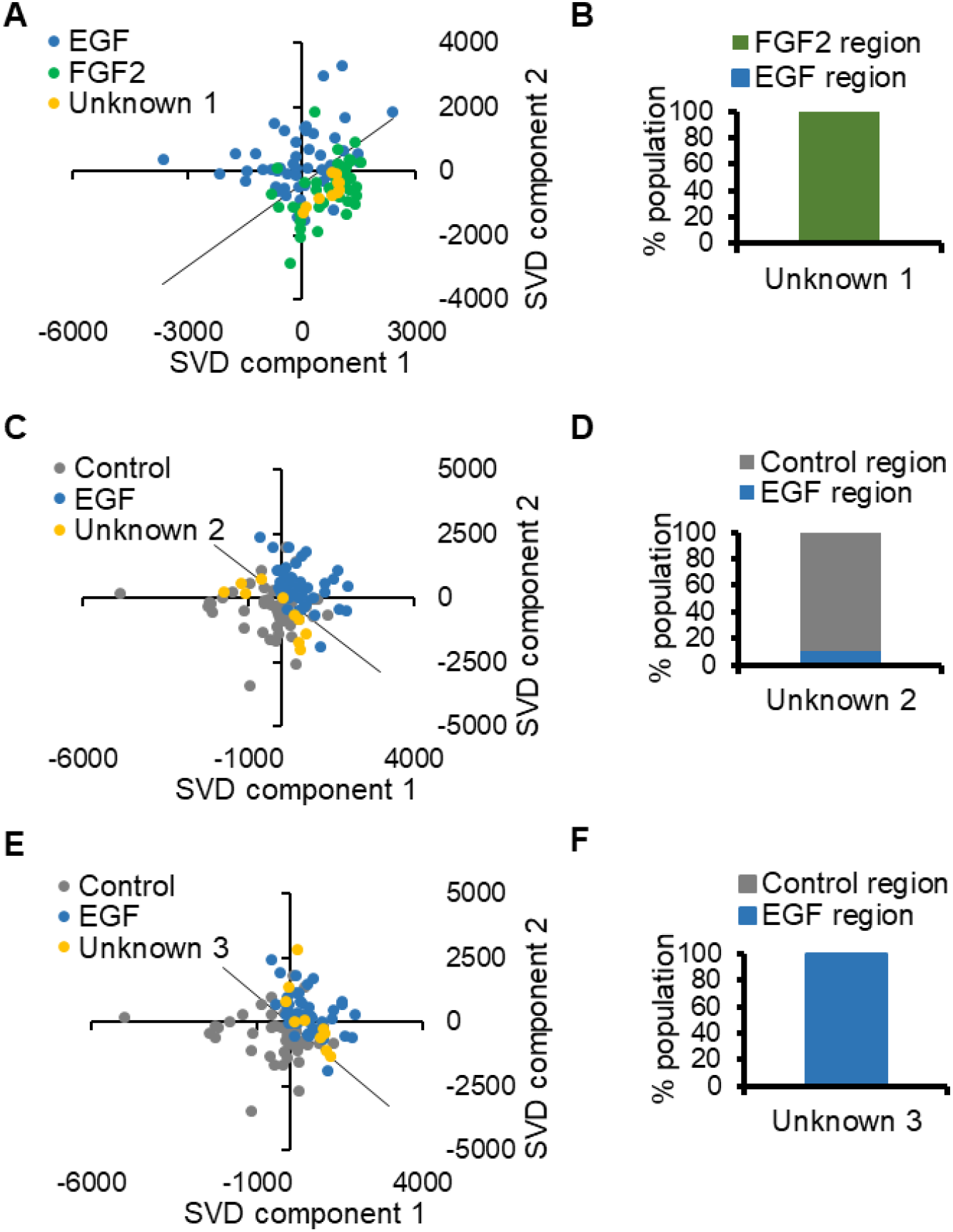
Unknown organoids identified using SVD analysis of Raman spectra. Raman spectra derived from the unknown (blinded) datasets of cell-dense organoid regions were subjected to the SVD analysis with known FGF2 (green), EGF (blue), and control (grey) datasets (derived from **Figure 3**). One representative plot is shown for each unknown, which demonstrates co-distribution with a known treatment. (**A**) SVD plot of unknown 1 with known EGF and FGF2 spectra. (**B**) % population of unknown 1 co-distributed with FGF2 or EGF. (**C**) SVD plot of unknown 2 with known EGF and control spectra. (**D**) % population of unknown 2 co-distributed with EGF or control. (**E**) SVD plot of unknown 3 with known EGF and control spectra. (**F**) % population of unknown 2 co-distributed with EGF or control (specific assignments of unknown samples detailed in **Table 2**). Additional SVD comparisons for unknowns 1-3 are provided in **Supplementary Figures 2-4**, respectively.

## Discussion

Organoid models are popular because they better represent the 3D cellular organization within an organ and the signaling between cells, for understanding organ development and for many translational applications [1,2]. However, the inherent heterogeneity in their development demands non-destructive approaches to verify the differentiation state or phenotype of the organoids. Here we report a Raman-based imaging method to circumvent challenges due to autofluorescence produced by the laminin-rich basement membrane that is commonly used in organoid cultures. We find that Raman micro-spectroscopy is ideal for organoid imaging, since the confocal microscope can be used to focus exclusively on cell-dense regions, with the out-of-focus light rejected, and post-processing can remove saturated Matrigel-rich spectra (**Figure 2C**). Unique Raman spectral signatures, which represent the average of multiple cellular regions, can then be defined for the desired developmental stage or phenotype (**Figure 3B**). We report that spectral signatures can be used to distinguish between organoids differentiated in the presence of FGF2 from those treated with EGF and from those not treated with a growth factor, by employing either multiple peak assignments or multivariate SVD analysis. Our results demonstrate the applicability of Raman spectral signatures for distinguishing between different phenotypes in the 3D context of the organoids.

In this study, we focused on the cell-dense regions of organoids, representing primarily epithelial cells, and the Raman signatures associated with each treatment (control, EGF, or FGF2). The differentiated FGF2 spectra had several significant Raman peaks that facilitated their separation from both control and EGF spectra. The differentiation and organization of the proacinar cells within the FGF2 samples produced the changes in the average spectra at broad peaks associated with Amide III, CH-deformation, and amide I (**Supplementary Figure 1**). This may be indicative of a differential distribution and organization of lipids and proteins in cells undergoing secretory proacinar differentiation. Since this approach is intended for the verification of differentiation into proacinar phenotype, and not for the identification of proacinar regions specifically, the cell-type specific Raman signatures were not assigned within this work. Importantly, Raman micro-spectroscopy facilitated the correct identification of organoid treatment using two separate methods of handling blinded datasets (**Figure 4-5**). Our results indicate that a higher throughput in identification and/or classification of organoids can be achieved by combining Raman imaging approach with multivariate data analysis, such as SVD. Ideally, this could be utilized in tracking the developmental stages of organoids and used to determine thresholds for production of more uniform samples.

Previously, Raman spectroscopy has been used to evaluate the mineral content of bone organoids matrices, to confirm scaffolding decellularization and nanomaterial for engineering of transplantable graphs [19–22], and to evaluate the drug response of sectioned, paraffin embedded tumor spheroids [23]. The utilization of Raman spectroscopy to differentiate between healthy and diseased tissue in numerous animal and human models has been well established, with far reaching clinical applications [6,8,9,24–26]. Our approach is novel as it focuses on unlabeled, non-destructive imaging of intact organoids, proposes a original method of rejecting the Matrigel dense signals, and defines a Raman signature for differentiated proacinar organoids. It could be converted into a high-throughput screening method to ensure conformity in organoid development without destruction of samples.

Machine learning has been applied in fluid samples for disease diagnostic approaches [27], suggesting its potential power for application in tissue- and organoid-based disease diagnostics. With further advances in image processing, Raman micro-spectroscopy combined with various data analysis techniques can be applied to many biological problems and therapeutic applications.

## Materials and Methods

### Mouse submandibular gland cell isolation

Embryonic day 16 (E16) timed-pregnant CD-1 female mice ordered from Charles River Laboratories were delivered to the University at Albany animal facility. First the E16 embryos and then the submandibular glands (SMGs) were removed using protocols approved by the University at Albany Institutional Animal Care and Use Committee (IACUC). SMG removal involved slicing the mandible with a scalpel and then removing the glands using sterile forceps under a dissecting microscope. Epithelial and mesenchymal cell populations were enriched, as described (Hossieni-1 and -2). Briefly, SMGs were microdissected in phosphate buffered saline (PBS) containing collagenase/hyaluronidase (StemCell Technologies, #7912) and dispase II (Life Technologies, #17105041) followed by manual trituration, and epithelial and mesenchymal cell populations were enriched by gravity sedimentation. The mesenchymal enriched cell population was further enriched by filtration through 70 µm (Falcon #087712) and 40 µm cell strainers (Fischer Scientific #22363547), washed by pelleting at 450xG for 5 minutes, and resuspended in media: Dulbecco’s Modified Eagle Medium/Nutrient Mixture F12 (DMEMF12, Fisher # 21041025) containing 5000 units/mL of penicillin and 5000 µg/mL of streptomycin (Pen-strep, Fisher # 15070063) and 10% fetal bovine serum.

### Epithelial organoid formation

About 900 epithelial clusters (1.0 gland equivalent) with a 20,000-50,000 stromal/mesenchymal cell addition (0.2 gland equivalent) were embedded in Matrigel (Corning #356234) at a 1:1 cell to Matrigel ratio. 10 µL of the cell and basement membrane mixture was seeded into the well of a 50 mm glass bottom dishes (MatTek #P50G-1.5-14F). The Matrigel was solidified by incubating at 37°C in a tissue culture incubator (Thermofisher Scientific Forma Series II) for 15 minutes, and covered with 180 µL of DMEM/F-12/10% FBS/PenStrep with or without growth factors added. The growth factor concentrations used were 100 ng/ml Epidermal Growth Factor (EGF) (PeproTech #AF100-15) or Fibroblast Growth Factor-2 (FGF2) (Peprotech #450-33) that were solubilized in 0.2% BSA and stored at -20°C in single-use aliquots. Organoids were cultured for 7 days at 37°C in a tissue culture incubator in 5% CO_2_ with the media replaced once at day 4. After 7 days of culturing, organoids were fixed by replacing the media with 4% paraformaldehyde (PFA) (Electron Microscopy Sciences #15710) in 1X PBS for 20 minutes and stored in 1X PBS prior to Raman imaging or immunocytochemistry.

### Immunocytochemistry, fluorescent imaging and analysis

Immunocytochemistry (ICC) was performed, as described previously [16,17], with 0.4% Triton-X 100 (Sigma #T9284-100ML) instead of 0.1% Triton-X for PFA-fixed samples. All primary antibody incubations were overnight at 4°C. Secondary antibody incubations were between 1-3 hours at room temperature. Primary antibodies and dilutions used include Aquaporin 5 (AQP5, 1:400; Alomone #AQP-005), Cytokeratin 7 (K7, 1:200; Abcam #ab9021), and epithelial cell adhesion molecule (EpCAM) directly conjugated to Fluorescein isothiocyanate (FITC) (1:400; eBiosciences #11-5791-82). Secondary antibodies, including Cyanine and Alexa dye-conjugated AffiniPure F(ab’)2 fragments, were purchased from Jackson ImmunoResearch Laboratories and used at a dilution of 1:250. 4′,6-diamidino-2-phenylindole (DAPI) (Life Technologies #D1306) was used for nuclei staining in conjunction with secondary antibodies. 100 µL of mounting media containing 90% (vol/vol) glycerol (Sigma #G5516-1L) in 1XPBS with 1-4 diazobicyclo[2,2,2]octane (DABCO) (Sigma #D27802-100G) and n-propyl gallate (Sigma #P3130-100G) as antifade agent was added directly on top of the organoids. Imaging was performed using a Zeiss Z1 Cell Observer widefield microscope or Zeiss LSM 710 confocal microscope at 10x, 20x or 63x (oil immersion) magnifications with the same configuration for all samples within an experiment. Quantification of immunostained area and organoid sizes was performed using FIJI v1.53c and post-processed with rolling-ball subtraction and line tool [18]. All statistical significances calculated between stained areas were done using a single-factor ANOVA followed with post-hoc Tukey HSD test for multi-sample comparison. Statistical tests were performed in Microsoft Excel (Microsoft Corporation) or R (R Core Team).

### Raman micro-spectroscopy collection and processing

Prior to Raman imaging, organoids were transferred to an imaging chamber (Warner) using quartz coverslips (Esco Optics). There organoids were inspected, and brightfield images collected on AmScope with a 4x objective. All Raman spectra were collected using a Horiba XploRA Plus Confocal Raman Microscope with a built-in 1024×256 TE air-cooled Syncerity CCD camera (pixel size 26 micron, temperature -60°C). A second camera, incorporated into the Horiba system, was employed for the collection of brightfield images of the samples on the stage. All Raman measurements of the organoids were made with a plan N 4x Olympus objective (Numerical Aperture (NA): 0.10, Infinity Corrected), 532 nm laser (100% full power), 1800 gr/mm grating, 500-2000 cm^-1^ spectral region, 100 µm slit, 300 µm confocal aperture, 30 s acquisition time and 3 accumulations per point, which were averaged by the program with an integrated spike removal feature during the active collection process.

To identify the cell dense regions within Matrigel, brightfield images of organoids were collected for determination of their X-Y coordinates, which was followed by point-by-point scans with multiple Z depths. Raman spectra collection of control, EGF-, and FGF2-treated salivary organoids for classification and blind studies focused on the cell-dense regions, with 5 locations scanned per image, with 3-4 different Z depths (step of 150 µm). A minimum of three organoids per treatment were evaluated within a dataset. After the processing and elimination of spectra which represented Matrigel-rich regions above or below the cell-dense target, 50 Raman spectra were reserved for classification of organoids, while 10 spectra per type were reserved for the blind study. Spectral processing was performed using the HORIBA LabSpec6 software, which included de-noise (first degree polynomial method with size 4), fluorescent background removal (second degree polynomial fit with 256 points), and normalization to the phenylalanine peak (∼ 1003 cm^-1^).

### Spectral analysis: peak ratios and singular value decomposition (SVD)

Peak ratio analysis was completed with a normalized spectral dataset in a spreadsheet (Excel, Microsoft). In-house Matlab code was used to implement the SVD analysis [10]. The Matlab code took stacked Raman spectra as the input n×m matrix, where n is the number of points in the spectrum and m is the number of Raman spectra in the dataset. The input matrix was decomposed into matrices U, Σ, and V^T^, where matrix V was used to generate SVD scatter plots, while individual SVD components were collected in the matrix U.

### Blinded evaluation of organoid spectra

The total of 10 spectra was excluded from each control, EGF, and FGF2 dataset and renamed unknown 1-3. The average peak intensity of unknown groups was acquired blind at 569, 621, 676, 1124, 1248, 1335, 1446, 1654, and 1927 cm^-1^ and compared to the known sample mean peaks; assignment of peaks was based on minimum difference of the means. Simple majority of individual peak assignments was used to determine the treatment type of unknown samples. The SVD process remained unchanged, except here the input matrices were generated by stacking different combinations of the Raman spectra. The first input matrix contained control Raman spectra (columns 1-50), EGF Raman spectra (columns 51-100) and unknown 1 (columns 101-110). The rows were the points of the spectral range, 500-2000 cm^-1^ (750 rows). The process was repeated for control, EGF and unknown 2; then control, EGF and unknown 3; control, FGF2 and unknown 1; control, FGF2 and unknown 2; control, FGF2 and unknown 3; EGF, FGF2 and unknown 1; EGF, FGF2 and unknown 2; and, finally, EGF, FGF2 and unknown 3.

## Abbreviations

AQP5: Aquaporin-5, used to identify proacinar cells
DAPI: 4′,6-diamidino-2-phenylindole, a nucleic acid dye
DMEM/F12: Dulbecco’s Modified Eagle Medium /Nutrient Mixture F12
EGF: Epidermal growth factor, used to promote growth not proacinar differentiation
EpCAM: Epithelial cell adhesion molecule, epithelial cell marker
FBS: fetal bovine serum
FGF2: Fibroblast growth factor 2, used to promote proacinar cell differentiation
FITC: Fluorescein isothiocyanate
K7: Keratin-7, used to identify ductal cells
Pen-Strep: penicillin and streptomycin

## Acknowledgments

We acknowledge the Life Science Research Building Core Facilities at University of Albany, State University of New York.

## Author contributions

Conceptualization: KT

Methodology: KT, NM, TK, DN, AS, ML, AK

Investigation: KT, NM, TK,

Visualization: KT, NM, TK,

Supervision: DN, AS, ML, AK

Writing—original draft: KT

Writing—review & editing: KT, NM, TK, DN, AS, ML, AK

## Data and materials availability

All data, code, and materials used in the analyses is available upon contacting the corresponding authors ML or AK

## Supplementary Materials

Please see attached Supplementary Materials

## References

[1] M. Hofer, M.P. Lutolf, Engineering organoids, Nature Reviews Materials. 6 (2021) 402–420. https://doi.org/10.1038/s41578-021-00279-y.

[2] Z.F. Hosseini, D.A. Nelson, N. Moskwa, M. Larsen, Generating Embryonic Salivary Gland Organoids, Current Protocols in Cell Biology. 83 (2019) 1–20. https://doi.org/10.1002/cpcb.76.

[3] A.C. Rios, H. Clevers, Imaging organoids: A bright future ahead, Nature Methods. 15 (2018) 24–26. https://doi.org/10.1038/nmeth.4537.

[4] I.A. Okkelman, N. Neto, D.B. Papkovsky, M.G. Monaghan, R.I. Dmitriev, A deeper understanding of intestinal organoid metabolism revealed by combining fluorescence lifetime imaging microscopy (FLIM) and extracellular flux analyses, Redox Biology. 30 (2020) 101420. https://doi.org/10.1016/j.redox.2019.101420.

[5] M. Yasunaga, Y. Fujita, R. Saito, M. Oshimura, Y. Nakajima, Continuous long-term cytotoxicity monitoring in 3D spheroids of beetle luciferase-expressing hepatocytes by nondestructive bioluminescence measurement, BMC Biotechnology. 17 (2017) 1–12. https://doi.org/10.1186/s12896-017-0374-1.

[6] Z. Movasaghi, S. Rehman, I.U. Rehman, Raman Spectroscopy of Biological Tissues, Applied Spectroscopy Reviews. 42 (2007) 493–541. https://doi.org/10.1080/05704920701551530.

[7] T. Huser, J. Chan, Raman spectroscopy for physiological investigations of tissues and cells, Advanced Drug Delivery Reviews. 89 (2015) 57–70. https://doi.org/10.1016/j.addr.2015.06.011.

[8] M.B. Fenn, P. Xanthopoulos, G. Pyrgiotakis, S.R. Grobmyer, P.M. Pardalos, L.L. Hench, Raman Spectroscopy for Clinical Oncology, Advances in Optical Technologies. 2011 (2011) 1–20. https://doi.org/10.1155/2011/213783.

[9] W.C. Zúñiga, V. Jones, S.M. Anderson, A. Echevarria, N.L. Miller, C. Stashko, D. Schmolze, P.D. Cha, R. Kothari, Y. Fong, M.C. Storrie-Lombardi, Raman Spectroscopy for Rapid Evaluation of Surgical Margins during Breast Cancer Lumpectomy, Scientific Reports. 9 (2019) 14639. https://doi.org/10.1038/s41598-019-51112-0.

[10] K. Tubbesing, T.C. Khoo, S. Bahreini Jangjoo, A. Sharikova, M. Barroso, A. Khmaladze, Iron-binding cellular profile of transferrin using label-free Raman hyperspectral imaging and singular value decomposition (SVD), Free Radical Biology and Medicine. 169 (2021) 416–424. https://doi.org/10.1016/j.freeradbiomed.2021.04.030.

[11] T.C. Khoo, K. Tubbesing, A. Rudkouskaya, S. Rajoria, A. Sharikova, M. Barroso, A. Khmaladze, Quantitative label-free imaging of iron-bound transferrin in breast cancer cells and tumors, Redox Biology. 36 (2020) 101617. https://doi.org/10.1016/j.redox.2020.101617.

[12] A. Khmaladze, J. Jasensky, E. Price, C. Zhang, A. Boughton, X. Han, E. Seeley, X. Liu, M.M.B. Holl, Z. Chen, Hyperspectral Imaging and Characterization of Live Cells by Broadband Coherent Anti-Stokes Raman Scattering (CARS) Microscopy with Singular Value Decomposition (SVD) Analysis, Applied Spectroscopy. 68 (2014) 1116–1122. https://doi.org/10.1366/13-07183.

[13] J. Jasensky, A.P. Boughton, A. Khmaladze, J. Ding, C. Zhang, J.E. Swain, G.W. Smith, Z. Chen, G.D. Smith, Live-cell quantification and comparison of mammalian oocyte cytosolic lipid content between species, during development, and in relation to body composition using nonlinear vibrational microscopy, Analyst. (2016). https://doi.org/10.1039/c6an00629a.

[14] R.J. Swain, M.M. Stevens, Raman microspectroscopy for non-invasive biochemical analysis of single cells, Biochemical Society Transactions. 35 (2007) 544–549. https://doi.org/10.1042/BST0350544.

[15] Z.F. Hosseini, D.A. Nelson, N. Moskwa, L.M. Sfakis, J. Castracane, M. Larsen, FGF2-dependent mesenchyme and laminin-111 are niche factors in salivary gland organoids, Journal of Cell Science. 131 (2018). https://doi.org/10.1242/jcs.208728.

[16] W.P. Daley, K.M. Gulfo, S.J. Sequeira, M. Larsen, Identification of a mechanochemical checkpoint and negative feedback loop regulating branching morphogenesis, Developmental Biology. 336 (2009) 169–182. https://doi.org/10.1016/j.ydbio.2009.09.037.

[17] S.J. Sequeira, D.A. Soscia, B. Oztan, A.P. Mosier, R. Jean-Gilles, A. Gadre, N.C. Cady, B. Yener, J. Castracane, M. Larsen, The regulation of focal adhesion complex formation and salivary gland epithelial cell organization by nanofibrous PLGA scaffolds, Biomaterials. 33 (2012) 3175–3186. https://doi.org/10.1016/j.biomaterials.2012.01.010.

[18] J. Schindelin, I. Arganda-Carreras, E. Frise, V. Kaynig, M. Longair, T. Pietzsch, S. Preibisch, C. Rueden, S. Saalfeld, B. Schmid, J.Y. Tinevez, D.J. White, V. Hartenstein, K. Eliceiri, P. Tomancak, A. Cardona, Fiji: An open-source platform for biological-image analysis, Nature Methods. 9 (2012) 676–682. https://doi.org/10.1038/nmeth.2019.

[19] A. Akiva, J. Melke, S. Ansari, N. Liv, R. Meijden, M. Erp, F. Zhao, M. Stout, W.H. Nijhuis, C. Heus, C. Muñiz Ortera, J. Fermie, J. Klumperman, K. Ito, N. Sommerdijk, S. Hofmann, An Organoid for Woven Bone, Advanced Functional Materials. 31 (2021) 2010524. https://doi.org/10.1002/adfm.202010524.

[20] L. Meran, I. Massie, S. Campinoti, A.E. Weston, R. Gaifulina, L. Tullie, P. Faull, M. Orford, A. Kucharska, A. Baulies, L. Novellasdemunt, N. Angelis, E. Hirst, J. König, A.M. Tedeschi, A.F. Pellegata, S. Eli, A.P. Snijders, L. Collinson, N. Thapar, G.M.H. Thomas, S. Eaton, P. Bonfanti, P. de Coppi, V.S.W. Li, Engineering transplantable jejunal mucosal grafts using patient-derived organoids from children with intestinal failure, Nature Medicine. 26 (2020) 1593–1601. https://doi.org/10.1038/s41591-020-1024-z.

[21] L. Sfakis, A. Sharikova, D. Tuschel, F.X. Costa, M. Larsen, A. Khmaladze, J. Castracane, Core/shell nanofiber characterization by Raman scanning microscopy, Biomedical Optics Express. 8 (2017) 1025. https://doi.org/10.1364/BOE.8.001025.

[22] A. Sharikova, Z.I. Foraida, L. Sfakis, L. Peerzada, M. Larsen, J. Castracane, A. Khmaladze, Characterization of nanofibers for tissue engineering: Chemical mapping by Confocal Raman microscopy, Spectrochimica Acta - Part A: Molecular and Biomolecular Spectroscopy. 227 (2020) 117670. https://doi.org/10.1016/j.saa.2019.117670.

[23] L.E. Jamieson, D.J. Harrison, C.J. Campbell, Raman spectroscopy investigation of biochemical changes in tumor spheroids with aging and after treatment with staurosporine, Journal of Biophotonics. 12 (2019). https://doi.org/10.1002/jbio.201800201.

[24] K. Sugiyama, J. Marzi, J. Alber, E.M. Brauchle, M. Ando, Y. Yamashiro, B. Ramkhelawon, K. Schenke-Layland, H. Yanagisawa, Raman microspectroscopy and Raman imaging reveal biomarkers specific for thoracic aortic aneurysms, Cell Reports Medicine. 2 (2021) 100261. https://doi.org/10.1016/j.xcrm.2021.100261.

[25] H.J. Butler, J.M. Cameron, C.A. Jenkins, G. Hithell, S. Hume, N.T. Hunt, M.J. Baker, Shining a light on clinical spectroscopy: Translation of diagnostic IR, 2D-IR and Raman spectroscopy towards the clinic, Clinical Spectroscopy. 1 (2019) 100003. https://doi.org/10.1016/j.clispe.2020.100003.

[26] A.Y.F. You, M.S. Bergholt, J.P. St-Pierre, W. Kit-Anan, I.J. Pence, A.H. Chester, M.H. Yacoub, S. Bertazzo, M.M. Stevens, Raman spectroscopy imaging reveals interplay between atherosclerosis and medial calcification in the human aorta, Science Advances. 3 (2017). https://doi.org/10.1126/sciadv.1701156.

[27] E. Ryzhikova, N.M. Ralbovsky, V. Sikirzhytski, O. Kazakov, L. Halamkova, J. Quinn, E.A. Zimmerman, I.K. Lednev, Raman spectroscopy and machine learning for biomedical applications: Alzheimer’s disease diagnosis based on the analysis of cerebrospinal fluid, Spectrochimica Acta - Part A: Molecular and Biomolecular Spectroscopy. 248 (2021) 119188. https://doi.org/10.1016/j.saa.2020.119188.

[28] N. Stone, C. Kendall, J. Smith, P. Crow, H. Barr, Raman spectroscopy for identification of epithelial cancers, Faraday Discussions. 126 (2004) 141–157. https://doi.org/10.1039/b304992b.

[29] C. Krafft, L. Neudert, T. Simat, R. Salzer, Near infrared Raman spectra of human brain lipids, Spectrochimica Acta - Part A: Molecular and Biomolecular Spectroscopy. 61 (2005) 1529–1535. https://doi.org/10.1016/j.saa.2004.11.017.

[30] G.J. Puppels, F.F.M. de Mul, C. Otto, J. Greve, M. Robert-Nicoudt, D.J. Arndt-Jovint, T.M. Jovin, Studying single living cells and chromosomes by confocal Raman microspectroscopy, Nature. 347 (1990) 301–303.

[31] Z. Huang, A. McWilliams, H. Lui, D.I. McLean, S. Lam, H. Zeng, Near-infrared Raman spectroscopy for optical diagnosis of lung cancer, International Journal of Cancer. 107 (2003) 1047–1052. https://doi.org/10.1002/ijc.11500.

[32] S. Clède, F. Lambert, C. Sandt, S. Kascakova, M. Unger, E. Harté, M.A. Plamont, R. Saint-Fort, A. Deniset-Besseau, Z. Gueroui, C. Hirschmugl, S. Lecomte, A. Dazzi, A. Vessières, C. Policar, Detection of an estrogen derivative in two breast cancer cell lines using a single core multimodal probe for imaging (SCoMPI) imaged by a panel of luminescent and vibrational techniques, Analyst. 138 (2013) 5627–5638. https://doi.org/10.1039/c3an00807j.

