## supplementary figures and tables for "Unlabeled salivary gland organoids have distinct Raman signatures following FGF2-induced proacinar cell differentiation"

**This PDF file includes:**

Supplementary Figures
Supplementary Figure 1. Raman Spectra with common broad peak assignments.
Supplementary Figure 2. Unknown 1- SVD comparisons with known datasets.
Supplementary Figure 3. Unknown 2- SVD comparisons with known datasets.
Supplementary Figure 4. Unknown 3- SVD comparisons with known datasets.

Supplementary Tables
Supplementary Table 1. Statistical summary for Figure 1C
Supplementary Table 2. Statistical summary for Figure 1D
Supplementary Table 3. Statistical summary for Figure 1E
Supplementary Table 4. Statistical summary for Figure 2A
Supplementary Table 5. Statistical summary for Figure 3C-K

**Supplementary Figure 1.** Raman Spectral graph from Figure 3B with common peak assignments based on molecular composition

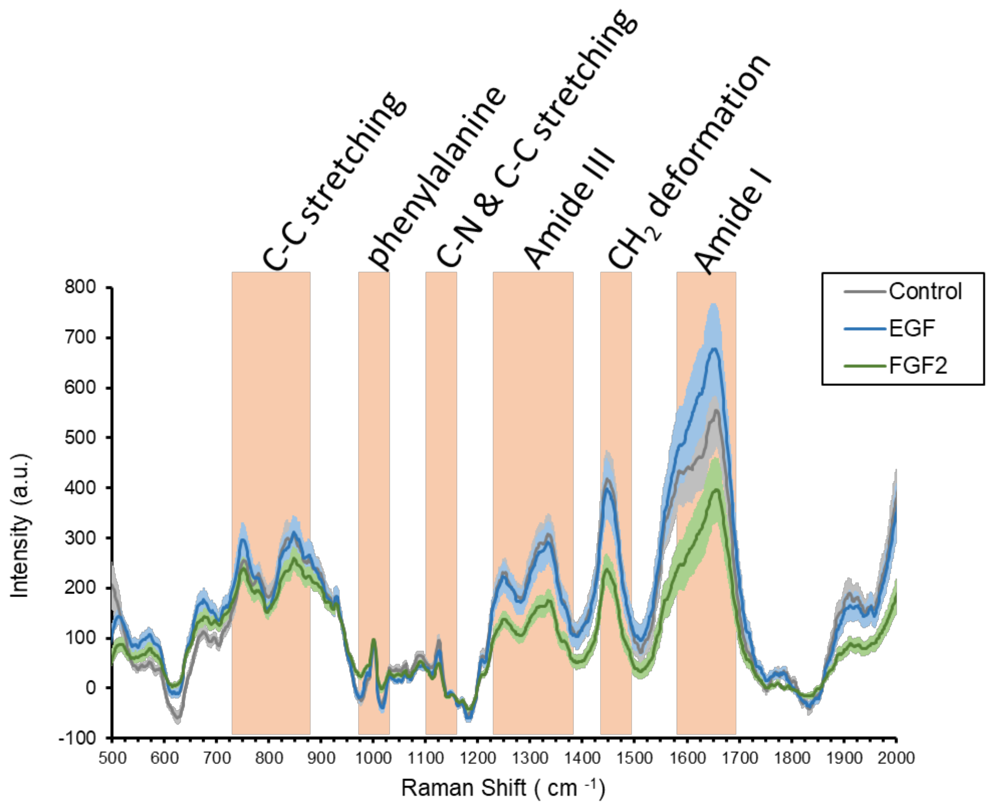


.

**Supplementary Figure 2.** Unknown 1- SVD comparisons with known datasets. For each comparison made there is the SVD scatter plot (A, C, D) and corresponding distribution of the unknown spectra based on scatter plot ( B, D, F).

*A-B are duplicated from Figure 5A and 5B.*

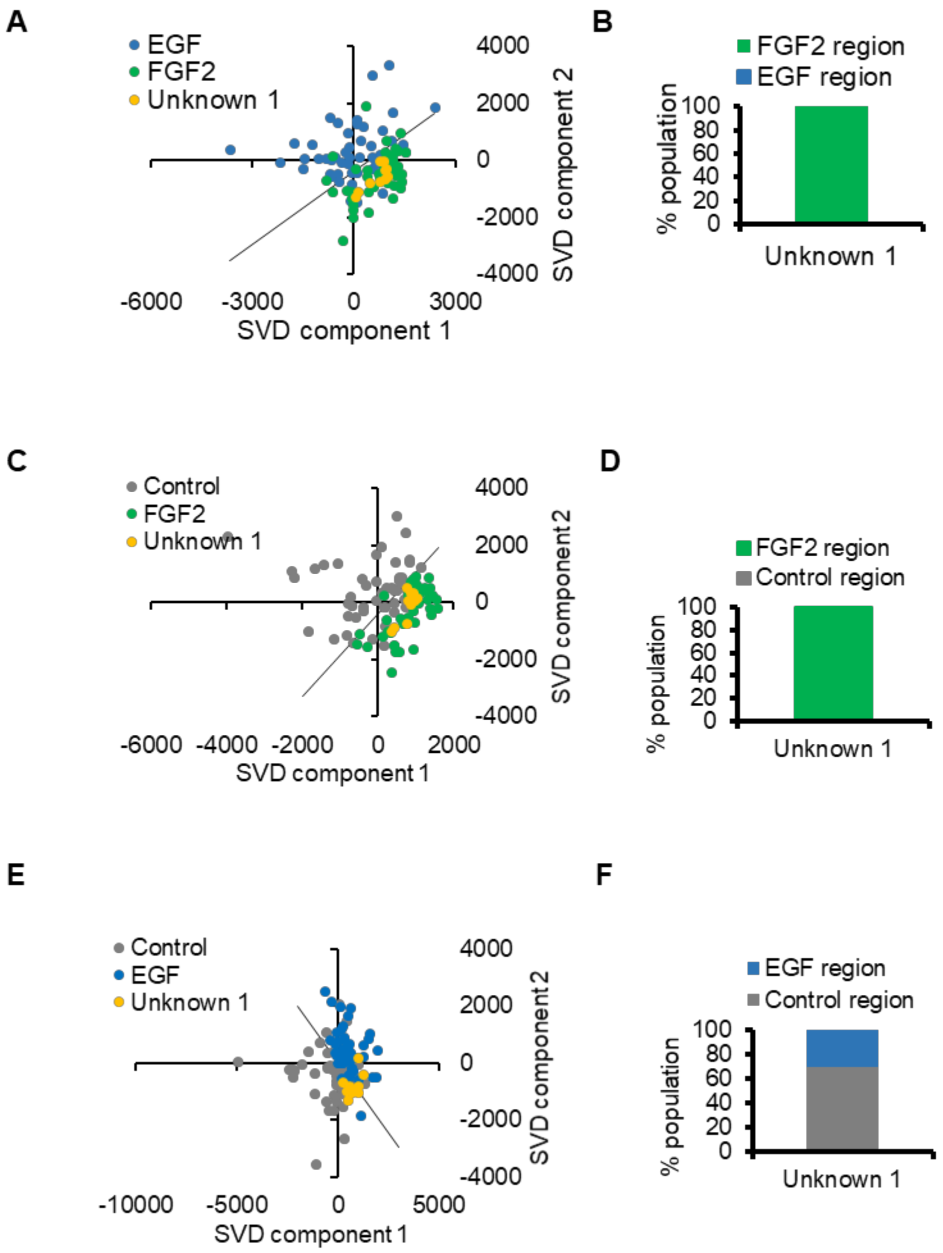

**Supplementary Figure 3.** Unknow 2 -SVD comparisons with known datasets. For each comparison made there is the SVD scatter plot (A, C, D) and corresponding distribution of the unknown spectra based on scatter plot ( B, D, F).

*A-B are duplicated from Figure 5C and 5D.*

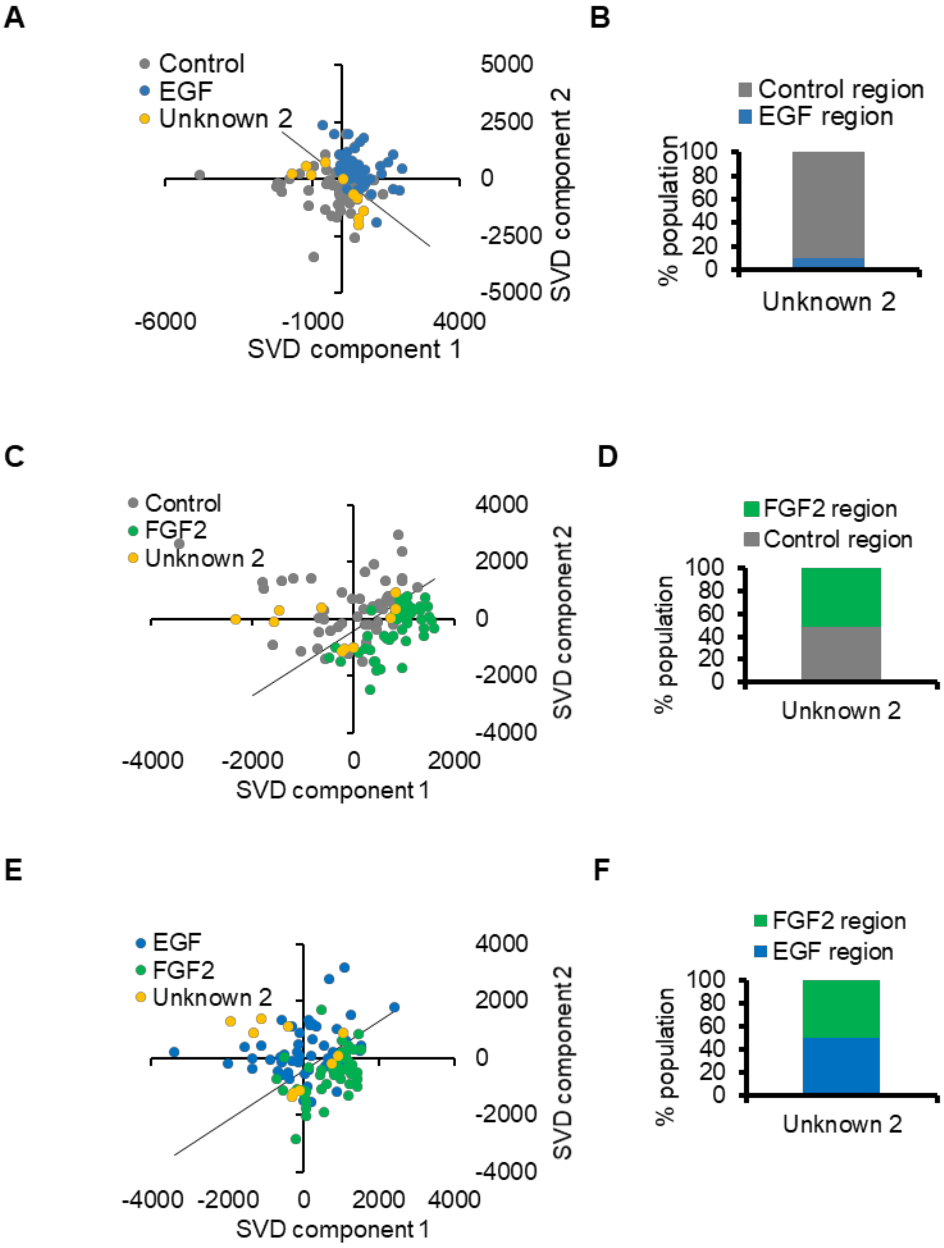

**Supplementary Figure 4.** Unknow 3 -SVD comparisons with known datasets. For each comparison made there is the SVD scatter plot (A, C, D) and corresponding distribution of the unknown spectra based on scatter plot ( B, D, F).

*A-B are duplicated from Figure 5E and 5F*

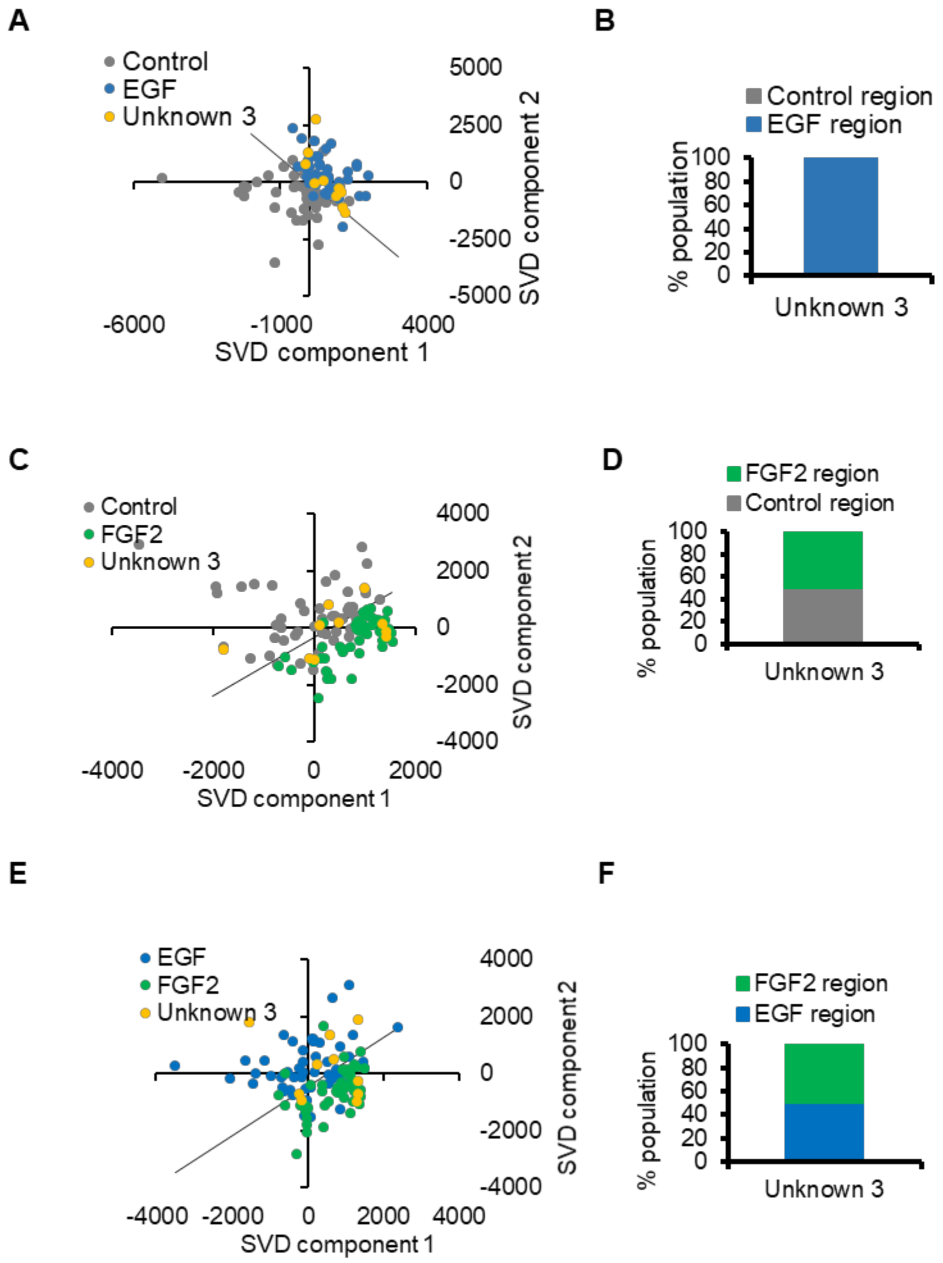

**Supplementary Table 1.**

| **Statistical summary for Figure 1C.** Asterisk indicates statistical significance. | | | | | |
|---|---|---|---|---|---|
| **SUMMARY** | | | | | |
| *Groups* | *Count* | *Sum* | *Average* | *Variance* | |
| Control | 3 | 0.13539 | 0.04513 | 0.00001 | |
| EGF | 3 | 0.06408 | 0.02136 | 0.00004 | |
| FGF2 | 3 | 0.96492 | 0.32164 | 0.00758 | |
| **ANOVA (single-factor)** | | | | | |
| *Source of Variation* | *SS* | *df* | *MS* | *P-value* | *F crit* |
| Between Groups | 0.16719 | 2 | 0.08360 | **0.00058*** | 5.1433 |
| Within Groups | 0.01526 | 6 | 0.00254 | | |
| Total | 0.18245 | 8 | | | |
| **Multiple comparison with Tukey HDS** Q-critical = 4.34 (Df =6, k=3, significance = 0.05) | | | | | |
| | Control vs. EGF | Control vs. FGF2 | EGF vs. FGF2 | | |
| HDS | 0.81642 | **9.49764*** | **10.31406*** | | |
| HDS> q-critical | No | **Yes*** | **Yes*** | | |
| Adj. p-value (anova-tukey in R) | 0.8368 | **0.0013*** | **0.0008*** | | |

**Supplementary Table 2.**

| **Statistical summary for Figure 1D.** Asterisk indicates statistical significance. | | | | | |
|---|---|---|---|---|---|
| **SUMMARY** | | | | | |
| *Groups* | *Count* | *Sum* | *Average* | *Variance* | |
| Control | 3 | 1.49317 | 0.49772 | 0.00968 | |
| EGF | 3 | 0.68749 | 0.22916 | 0.00385 | |
| FGF2 | 3 | 0.60166 | 0.20055 | 0.00467 | |
| **ANOVA (single-factor)** | | | | | |
| *Source of Variation* | *SS* | *df* | *MS* | *P-value* | *F crit* |
| Between Groups | 0.16125 | 2 | 0.08063 | **0.00624*** | 5.14325 |
| Within Groups | 0.03639 | 6 | 0.00606 | | |
| Total | 0.19764 | 8 | | | |
| **Multiple comparison with Tukey HDS** Q-critical = 4.34 (Df =6, k=3, significance = 0.05) | | | | | |
| | *Control vs. EGF* | *Control vs. FGF2* | *EGF vs. FGF2* | | |
| HDS | **5.97315*** | **6.60947*** | 0.63632 | | |
| HDS> q-critical | **Yes*** | **Yes*** | no | | |
| Adj. p-value (anova-tukey in R) | **0.0131*** | **0.0082*** | 0.8963 | | |

**Supplementary Table 3.**

| **Statistical summary for Figure 1E.** Asterisk indicates statistical significance. | | | | | |
|---|---|---|---|---|---|
| **SUMMARY** | | | | | |
| *Groups* | *Count* | *Sum* | *Average* | *Variance* | |
| Control | 3 | 0.281071 | 0.09369 | 0.000591 | |
| EGF | 3 | 0.278383 | 0.092794 | 2.51E-05 | |
| FGF2 | 3 | 4.88074 | 1.626913 | 0.01137 | |
| **ANOVA (single-factor)** | | | | | |
| *Source of Variation* | *SS* | *df* | *MS* | *P-value* | *F crit* |
| Between Groups | 4.704294 | 2 | 2.352147 | **1.3E-07** | 5.143253 |
| Within Groups | 0.023972 | 6 | 0.003995 | | |
| Total | 4.728265 | 8 | | | |
| **Multiple comparison with Tukey HDS** Q-critical = 4.34 (Df =6, k=3, significance = 0.05) | | | | | |
| | Control vs. EGF | Control vs. FGF2 | EGF vs. FGF2 | | |
| HDS | 0.0245 | **42.0138*** | **42.0383*** | | |
| HDS> Q-critical | no | **Yes*** | **Yes*** | | |
| Adj. p-value (anova-tukey in R) | 0.9998 | **3.94E-7*** | **3.93E-7*** | | |

**Supplementary Table 4.**

| **Statistical summary for Figure 2A.** Asterisk indicates statistical significance. | | | | |
|---|---|---|---|---|
| **SUMMARY** | | | | |
| *Groups* | *Count* | *Sum* | *Average* | *Variance* |
| Control | 10 | 41723.13 | 4172.31 | 2217675.07 |
| EGF | 7 | 49105.25 | 7015.04 | 739248.65 |
| FGF2 | 10 | 79305.33 | 7930.53 | 3993123.19 |

| **ANOVA (single-factor)** | | | | | |
|---|---|---|---|---|---|
| *Source of Variation* | *SS* | *df* | *MS* | *P-value* | *F crit* |
| Between Groups | 75435807 | 2 | 37717903 | **5.93E-5** | 3.4028 |
| Within Groups | 60332676 | 24 | 2513861 | | |
| Total | 135768483 | 26 | | | |

**Multiple comparison with Tukey HDS**
Q-critical = 3.532 (Df =24, k=3, significance = 0.05)

| | *Control vs. EGF* | *Control vs. FGF2* | *EGF vs. FGF2* |
|---|---|---|---|
| HDS | **4.7437*** | **7.4957*** | 1.5277 |
| HDS> q-critical | **Yes*** | **Yes*** | no |
| Adj. p-value (anova-tukey in R) | **0.0036** | **5.6E-5*** | 0.4810 |

### Supplementary Table 5.

| **Statistical summary for Figure 2A.** Bold with Asterisk indicates statistical significance for anova, while significant comparisons are listed for post hoc and additional | | | | |
|---|---|---|---|---|
| | *Anova, single-factor, with Dunnetts Post hoc test Used to determine significance reported on graphs* | | | |
| *Wavenumber ( $cm^{-1}$)* | *p-value-between groups* | *F value (F crit = 3.06)* | *dunnett's critical value-significant comparisons listed* | *Significant p-values listed 2 tailed, Student t-test (0.05, Bonferroni correction)* |
| 569 | **6.55E-08*** | 18.55 | **Control vs. EGF***<br>**Control vs. FGF2***<br>**EGF vs. FGF2*** | Control vs. EGF= 7.62E-09<br>Control vs. FGF2=1.04E-03<br>EGF vs. FGF2=7.76E-03 |
| 621 | **2.47E-14*** | 39.07 | **Control vs. EGF***<br>**Control vs. FGF2*** | Control vs. EGF= 4.01E-07<br>Control vs. FGF2= 5.30E-14<br>EGF vs. FGF2= 9.81E-03 |
| 675 | **1.29E-05*** | 12.17 | **Control vs. EGF***<br>**EGF vs. FGF2*** | Control vs. EGF= 8.06E-06<br>Control vs. FGF2= 1.05E-02<br>EGF vs. FGF2= 1.42E-02 |
| 1124 | **1.36E-09*** | 23.53 | **Control vs. EGF***<br>**Control vs. FGF2***<br>**EGF vs. FGF2*** | Control vs. EGF= 7.08E-03<br>Control vs. FGF2= 1.00E-09<br>EGF vs. FGF2= 8.94E-06 |
| 1248 | **5.28E-07*** | 15.97 | **Control vs. FGF2***<br>**EGF vs. FGF2*** | Control vs. FGF2= 1.66E-07<br>EGF vs. FGF2= 4.11E-06 |
| 1335 | **8.97E-07*** | 15.33 | **Control vs. FGF2***<br>**EGF vs. FGF2*** | Control vs. FGF2= 3.53E-07<br>EGF vs. FGF2= 3.99E-06 |
| 1446 | **2.30E-06*** | 14.20 | **Control vs. FGF2***<br>**EGF vs. FGF2*** | Control vs. FGF2= 5.40E-07<br>EGF vs. FGF2= 9.85E-06 |
| 1654 | **5.36E-06*** | 13.20 | **Control vs. FGF2***<br>**EGF vs. FGF2*** | Control vs. FGF2= 1.58E-03<br>EGF vs. FGF2= 3.39E-06 |
| 1927 | **3.22E-07*** | 16.58 | **Control vs. FGF2***<br>**EGF vs. FGF2*** | Control vs. FGF2= 9.73E-08<br>EGF vs. FGF2= 2.98E-06 |
